# Autofluorescence imaging reveals the impact of cryopreservation on T cell metabolism and activation response

**DOI:** 10.1101/2025.11.14.688556

**Authors:** Dan L. Pham, Meghana Kalluri, Cole Weaver, Angela Hsu, Amani Gillette, Wenxuan Zhao, Tyce Kearl, Peiman Hematti, Nirav Shah, Melissa C. Skala

## Abstract

Cryopreservation or the process of freezing cells is a cornerstone of most cell therapy protocols. Optimization of cryopreservation protocols and cryoprotectant agents to improve cell viability and functionality is under further investigation. However, the impact of cryopreservation on cellular metabolism and function immediately post-thaw is not fully understood. Here, we used label-free, non-invasive optical metabolic imaging (OMI) of NAD(P)H and FAD to characterize the activation response of frozen T cells from healthy donors and lymphoma patients post-thaw. Using OMI, we identified significant metabolic shift, along with delayed and diminished activation response in healthy donor T cells throughout the first 4.5 hours upon thawing. In cryopreserved peripheral T cells from lymphoma patients in our bispecific CD19/CD20 CAR T therapy clinical trial, OMI could identify early metabolic stress and allowed gating of metabolically-fit cells associated with post-thaw viability. Notably, in our pilot study, only metabolically-fit T cells from complete responders exhibited metabolic responses to activating stimuli within the first 4.5 hours post-thaw. Overall, our findings suggest that 4-5 hours post-thaw is a critical time window to assess the impact of cryopreservation and thawing, and support the potential of OMI to optimize cryopreservation protocols and evaluate patient T cell quality for cell therapy.

## INTRODUCTION

Cryopreservation is commonly used for long-term preservation of cells, tissue and other biological samples, which involves using low temperature (-80°C to -196°C) to slow down cell metabolism and other functional activity. Cryopreservation plays a central role for several cell therapies including Chimeric Antigen Receptor (CAR) or engineered T cell receptor (TCR) therapy. The complex manufacturing process of these cell therapies requires specialized facilities and expertise, and often employs a central manufacturing model that relies on cryopreservation of both the starting materials and final products^1^. For instance, current commercial CAR T manufacturing workflows start with collection of leukapheresis from the patient at the hospital site as the starting material. Patient’s leukapheresis products are transported to the manufacturing facility, where T cells are activated, transduced to express CAR transgene, and expanded^2^. At the end of the manufacturing process, the CAR T cell product is cryopreserved. This allows for preservation of the product while necessary quality control testing is performed, and for transportation to the hospital for infusion into the patient. Despite ongoing research to define the optimal cryopreservation protocol such as cryoprotectant agents, cooling rate, and post-thaw condition, the successful recovery of cell viability and function upon thawing remains challenging^2, 3^. Meanwhile, due to the toxic effect of residual cryoprotectant agent, such as dimethyl sulfide (DMSO), on immune cell viability and effector functions^4^, most CAR T cell products mandate infusion within 30-90 minutes after thawing^5^. Since proper T cell function is critical for the efficacy of CAR T cell therapy, it is important to characterize the impact of cryopreservation on T cells within a short time frame post-thaw, particularly concerning T cell activation response, a key function for CAR T manufacturing success and clinical outcome.

During CAR T cell manufacturing, T cell activation is induced by targeting stimulatory and co-stimulatory receptors on the T cell surface, such as CD3, CD28, and CD2^6^. Activation primes T cells for *CAR* gene transfer while followed by a proliferation phase to achieve a sufficient CAR T cell dosage, both of which are required for successful CAR T cell manufacturing^7^. Most CAR T products are frozen while going under quality control testing before their release. Antigen-specific activation of CAR T cells upon thawing and infusion into the patient and interaction with target tumor cells initiates cytotoxic functions to eliminate cancer cells, leading to tumor regression and patient treatment response. Therefore, understanding how cryopreservation affects CD3-mediated and antigen-specific activation of T cells upon thawing will allow optimization of cryopreservation protocols that ultimately enhance CAR T cell performance.

For T cell activation, metabolic reprogramming is crucial, not only to provide the necessary energy and molecules for effector function, but also to shape the epigenetic landscape that determines T cell differentiation fate^8–10^. Cryopreservation has been shown to affect cell viability and metabolism-related proteins while reducing metabolic activity in various mammalian single cell or multi-cellular models such as embryos and other reproductive cells^11,12,13^, engineered tissues (osteoblasts^14^), and transplanted hepatocytes^15,16^ or pancreatic islets^17,18^. However, the impact of cryopreservation is cell-type dependent^19^, and immediate metabolic changes in T cells post-thaw and their implication on early activation response remain unclear. We propose continuous monitoring of T cell metabolism in real time throughout a short window immediately post-thaw and in response to activation stimuli could provide essential insight for future improvements in the freeze-thaw processes.

While several bioassays exist for metabolism and T cell activation assessments, most of them either require destructive and time-consuming sample preparation that is not suitable for frequent analysis of the same culture over time, or do not offer single-cell resolution to characterize heterogeneous cell function. To address these challenges, we have developed optical metabolic imaging (OMI), a label-free non-invasive imaging method that allows assessment of single cell metabolism based on autofluorescent signals from metabolic coenzymes NAD(P)H and FAD^20,21,22^. NAD(P)H and FAD are an electron donor and acceptor, respectively, and are essential components of many metabolic pathways, including those important for T cell function and activation response such as glycolysis^23,24^. OMI yields 13 cellular metabolic features based on fluorescence intensity and lifetime of NAD(P)H and FAD (NAD(P)H intensity, τ_m_, τ_1_, τ_2_, α_1_, α_2_, FAD intensity, τ_m_, τ_1_, τ_2_, α_1_, α_2_, optical redox ratio (ORR)) and cell size. ORR is defined as the ratio of NAD(P)H intensity over the sum of NAD(P)H intensity and FAD intensity, which has been correlated to the cellular redox balance^25,26^. Meanwhile, fluorescence lifetime is defined as time taken for a molecule in the excited state to decay back to the ground state and emit a fluorescence signal. Free NAD(P)H self-quenches, resulting in a short fluorescence lifetime of around 400ps^27,28^. Upon protein-binding, NAD(P)H undergoes conformational changes, leading to long fluorescence lifetime of around 2.2-2.5ns^28,29^. FAD displays a reversed trend in fluorescence lifetime, with free FAD having a long lifetime and bound FAD having a short lifetime^24,30^. Therefore, the fluorescence lifetimes of NAD(P)H and FAD are indicative of their binding activity.

Previously, we demonstrated that metabolic imaging based on autofluorescence from NAD(P)H and FAD is sensitive and specific in characterizing metabolic changes upon T cell activation^31,32^. Here, we used OMI to determine the impact of cryopreservation on T cell metabolism and activation response, along with implications of cryopreservation for adoptive T cell therapy. Using OMI, we also evaluated the metabolic response accompanying different modes of T cell activation, including CD3-driven and antigen-specific activation in cryopreserved cells.

## RESULTS

### OMI identifies significant changes in frozen T cell metabolism from healthy donors upon thawing

CD3 T cells were isolated from peripheral blood of 3 healthy donors and divided in half for cryopreservation and fresh culture for 24 hours. Upon thawing, cryopreserved and fresh T cells from matched donor were imaged simultaneously using OMI every hour (**Fig 1A**). Throughout the 4-hour imaging time course, fresh quiescent T cells displayed stable metabolism, with no significant changes in NAD(P)H mean lifetime (NAD(P)H τ_m_) and a slight increase in the proportion of free NAD(P)H (NAD(P)H α_1_) at the 4.5-hour time point (**Fig 1B, C**). Interestingly, we observed significant and consistent metabolic changes in frozen quiescent T cells upon thawing (**Fig 1D, E**). Across 3 donors, frozen T cells showed a gradual but significant increase in NAD(P)H τ_m_ and decrease in NAD(P)H α_1_ within the first 4.5 hours post-thaw. To determine the effect sizes of metabolic changes over time, we calculated Glass’s Δ at each time point with respect to the 0.5-hour time point (**Fig 1F, G**). Glass’s Δ revealed a small effect of imaging time on fresh quiescent T cell metabolism. For several OMI parameters, the average effect size across 3 donors at each time point remain within -0.1 to 0.1, further indicating stable metabolism (**Fig 1F, G -left**). Meanwhile, we observed a large effect of time post-thaw on frozen quiescent T cell metabolism. Over time, Glass’s Δ for several OMI parameters of frozen T cells strengthened and reached an average value of greater than 1 (for NAD(P)H τ_m_) or smaller than -1 (for NAD(P)H α_1_) at the 4.5-hour time point (**Fig 1F, G -right**). This indicated that the differences in these OMI parameters exceeded one standard deviation throughout the imaging time course. Using OMI, we have determined consistent and significant metabolic changes in frozen T cells, with a shift towards increasing NAD(P)H τ_m_ and decreasing NAD(P)H α_1_ that continued up to 4.5-hour post thaw.

**Figure 1.**
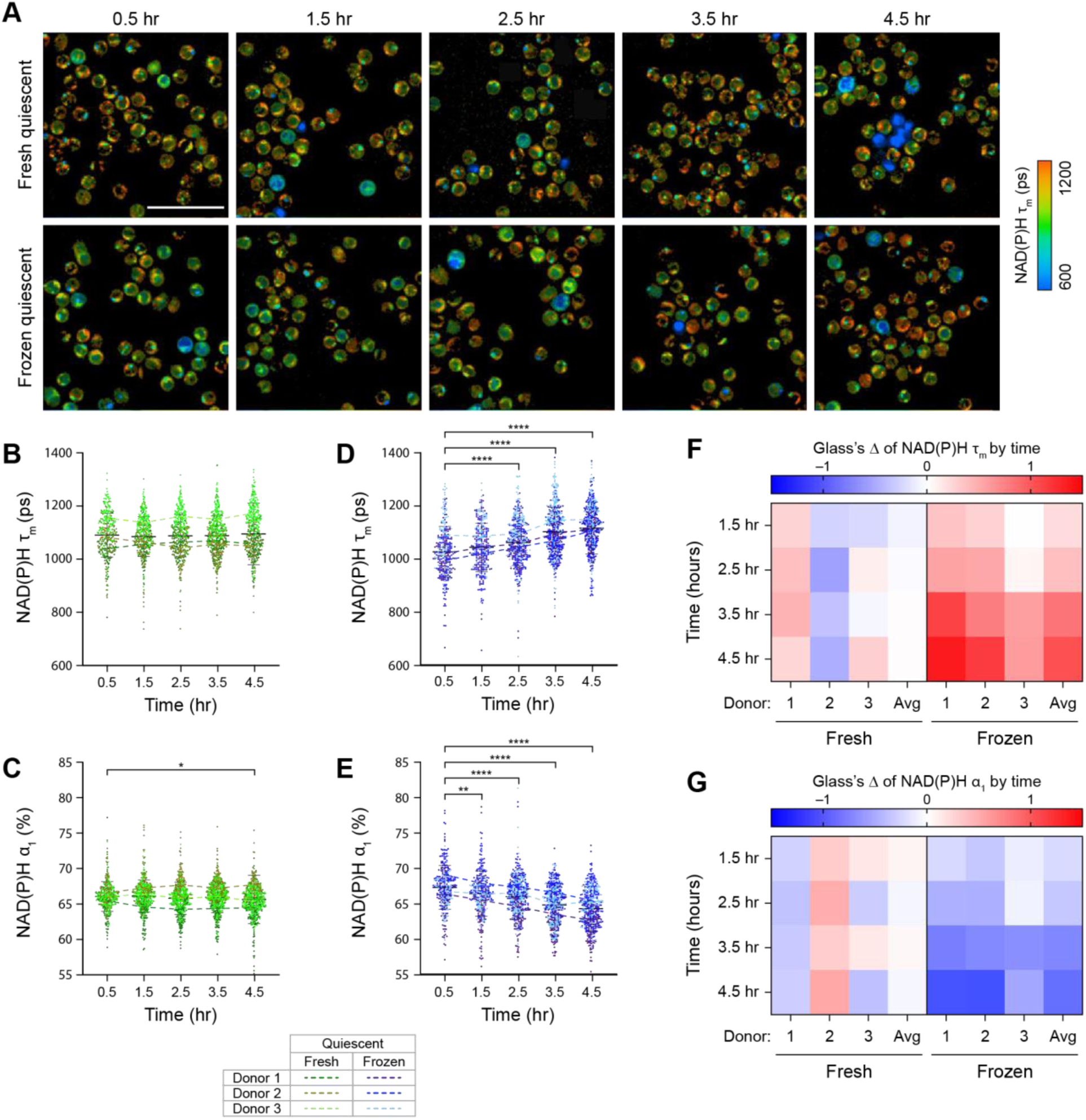
Cryopreserved T cells underwent significant metabolic changes upon thawing. **(A)** Representative NAD(P)H mean lifetime (NAD(P)H τ_m_) images of donor-matched fresh (top) and frozen (bottom) CD3 T cells throughout a 4.5-hour imaging time course. **(B-E)** Quantification of NAD(P)H τ_m_ and free NAD(P)H proportion (NAD(P)H α_1_) of fresh **(B-C)** or frozen **(D-E)** T cells. n = 299-537 cells/condition/time point across 3 biologically independent donors. Dots represent individual cells, color coded by condition (fresh or frozen) and donor. Two-sided non-parametric Kruskal-Wallis test with Dunn’s post hoc test for multiple comparison against corresponding OMI measurements at the 0.5-hour time point. **(F-G)** Glass’s Δs to quantify effect sizes of changes over time in **(F)** NAD(P)H τ_m_ and **(E)** NAD(P)H α_1_ of fresh and frozen T cells, with respect to the corresponding OMI measurements at the 0.5-hour time point. Glass’s Δ was reported for individual donors and the average values across 3 donors. Left 4 columns: fresh T cells, right 4 columns: frozen T cells. Scale bar is 50μm. Bars are mean ± standard deviation. * p < 0.05, ** p < 0.01, **** p < 0.0001.

### Fresh and frozen T cells from healthy donors displayed distinct metabolic response to activation during the first 4.5 hours

Due to significant changes in frozen T cell metabolism upon thawing, we further investigated how cryopreservation impacts T cell activation response using OMI. Frozen T cells from 3 healthy donors were immediately activated with StemCell αCD2/αCD3/αCD28 upon thawing. Similarly, donor-matched fresh T cells were also activated at the same time (**Fig 2A**). In fresh T cells, we observed the characteristic metabolic changes, with low NAD(P)H τ_m_ and high NAD(P)H α_1_, following activation (**Fig 2B, C**). This is consistent with previous studies showing a shift towards glycolysis in activated T cells^8,9^. These changes in T cell metabolism occurred early and were significant within 1 hour of activation. Activation-induced metabolic changes in fresh T cells continued to progress, with decreasing NAD(P)H τ_m_ and increasing NAD(P)H α_1_ observed over time (**Fig 2B, C**). We did not, however, observe similar metabolic changes in frozen activated T cells (**Fig 2D, E**). Across 3 donors, frozen activated T cells showed increased NAD(P)H τ_m_ and decreased NAD(P)H α_1_ that were sustained throughout the 4-hour imaging time course. These metabolic changes were opposite of donor-matched fresh activated T cells (**Fig 2B, C**) but were consistent with frozen quiescent T cells (**Fig 1**).

**Figure 2.**
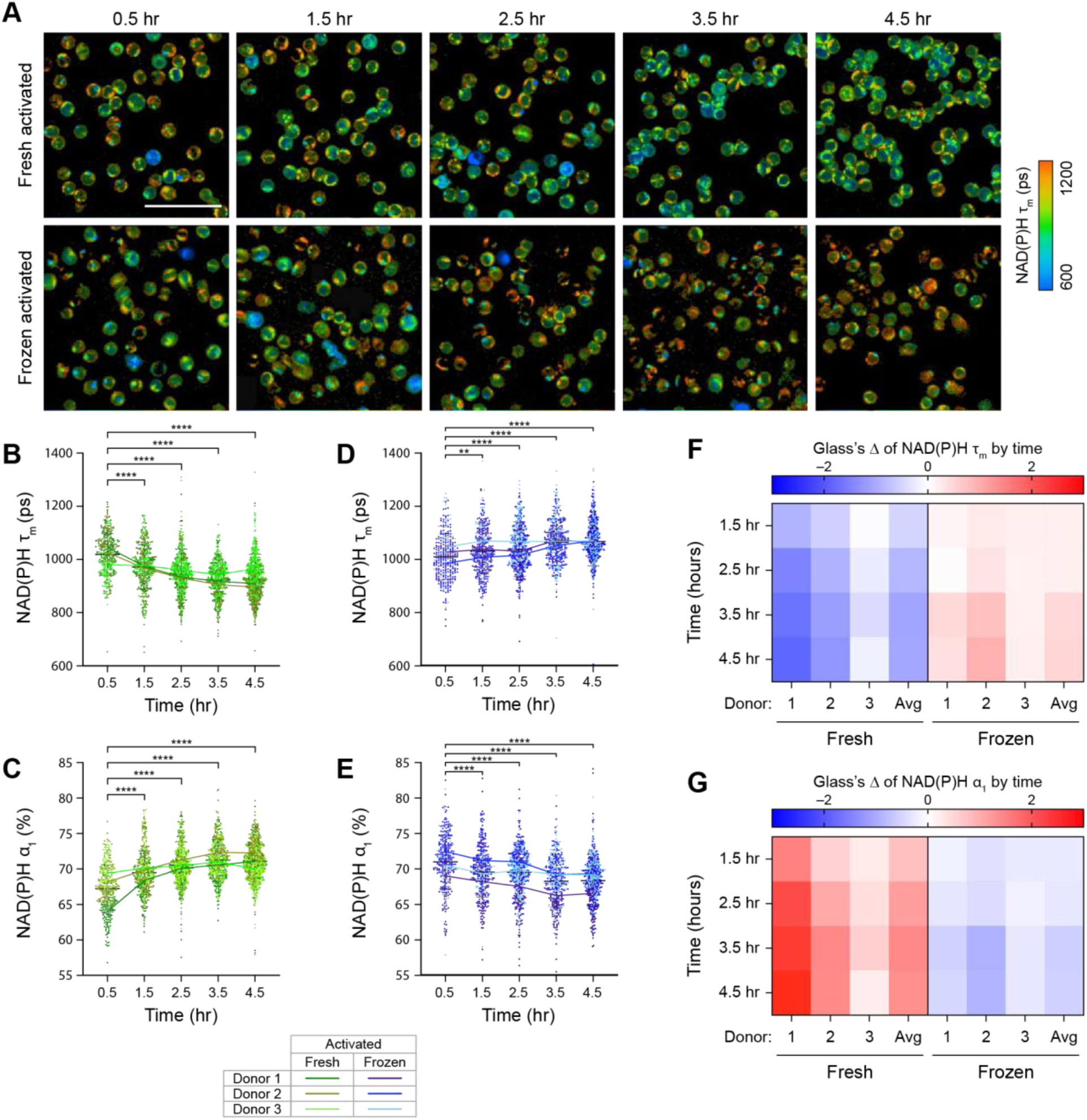
Fresh and frozen T cells displayed distinct metabolic response towards activating stimulus. **(A)** Representative NAD(P)H τ_m_ images of donor-matched fresh (top) and frozen (bottom) T cells upon activation. Frozen T cells were activated immediately after thawing. **(B-E)** Quantification of NAD(P)H τ_m_ and NAD(P)H α_1_ of **(B-C)** fresh or **(D-E)** frozen activated T cells. n = 296-618 cells/condition/time point across 3 biologically independent donors. Two-sided non-parametric Kruskal-Wallis test with Dunn’s post hoc test for multiple comparison against corresponding OMI measurements at the 0.5-hour time point. **(F-G)** Glass’s Δs to quantify effect sizes of changes over time in **(F)** NAD(P)H τ_m_ and **(E)** NAD(P)H α_1_ of fresh and frozen activated T cells, with respect to the corresponding OMI measurements at the 0.5-hour time point. Glass’s Δ was reported for individual donors and the average values across 3 donors. Left 4 columns: fresh T cells, right 4 columns: frozen T cells. Scale bar is 50μm. Bars are mean ± standard deviation. * p < 0.05, ** p < 0.01, **** p < 0.0001.

Glass’s Δ with respect to the 0.5-hour time point further confirmed opposite metabolic responses of donor-matched activated fresh and frozen T cells over time (**Fig 2F, G**). Throughout activation (0.5- to 4.5-hour), fresh T cells showed large effects in NAD(P)H τ_m_ and NAD(P)H α_1_ that strengthened over time, with average absolute Glass’s Δ values exceeding 1.2 at the 4.5-hour time point (**Fig 2F, G – left**). These findings demonstrated that fresh T cell metabolism, especially NAD(P)H binding activity, were highly responsive to activating stimulus. Meanwhile, for frozen T cells, we observed weak- to moderate-effect sizes throughout the imaging time course (**Fig 2F, G – right**). Interestingly, at 4.5 hours, average Glass’s Δ of frozen activated T cells (0.60 for NAD(P)H τ_m_ and -0.68 for NAD(P)H α_1_) were smaller than those of frozen quiescent cells (1.03 for NAD(P)H τ_m_ and -1.01 for NAD(P)H α_1_, respectively). We hypothesized that since thawing (increased NAD(P)H τ_m_ and decreased NAD(P)H α_1_) and activation (decreased NAD(P)H τ_m_ and increased NAD(P)H α_1_) induced opposite changes in NAD(P)H lifetime parameters, the effects on frozen activated T cells were confounded. However, since we observed an overall similar metabolic change (increased NAD(P)H τ_m_ and decreased NAD(P)H α_1_) in both frozen quiescent and frozen activated T cells within the first 4.5-hour post-thaw, our findings suggested that the effect of cryopreservation/thawing was dominant during this time window, and this potentially affected the ability of frozen T cells to respond to activating stimulus.

### OMI revealed delayed activation response in frozen T cells post-thaw

Since we observed different metabolic changes in fresh and frozen T cells upon activation, we further characterized the activation response in these groups compared to their donor-matched quiescent groups. We observed significantly lower NAD(P)H τ_m_ and higher NAD(P)H α_1_ between quiescent and activated fresh T cells as early as 0.5 hour after activation. The differences in these NAD(P)H lifetime parameters between fresh quiescent and fresh activated T cells further widened as the activation duration increased (**Fig 3A, B**). We also observed significantly higher normalized NAD(P)H intensity in fresh activated T cells compared to donor-matched fresh quiescent cells (**Supplementary Fig 2A**). The increase in NAD(P)H intensity, decrease in NAD(P)H τ_m_, and increased in NAD(P)H α_1_ indicated an increase in free NAD(P)H abundance following activation, consistent with previous studies ^33,34^.

**Figure 3.**
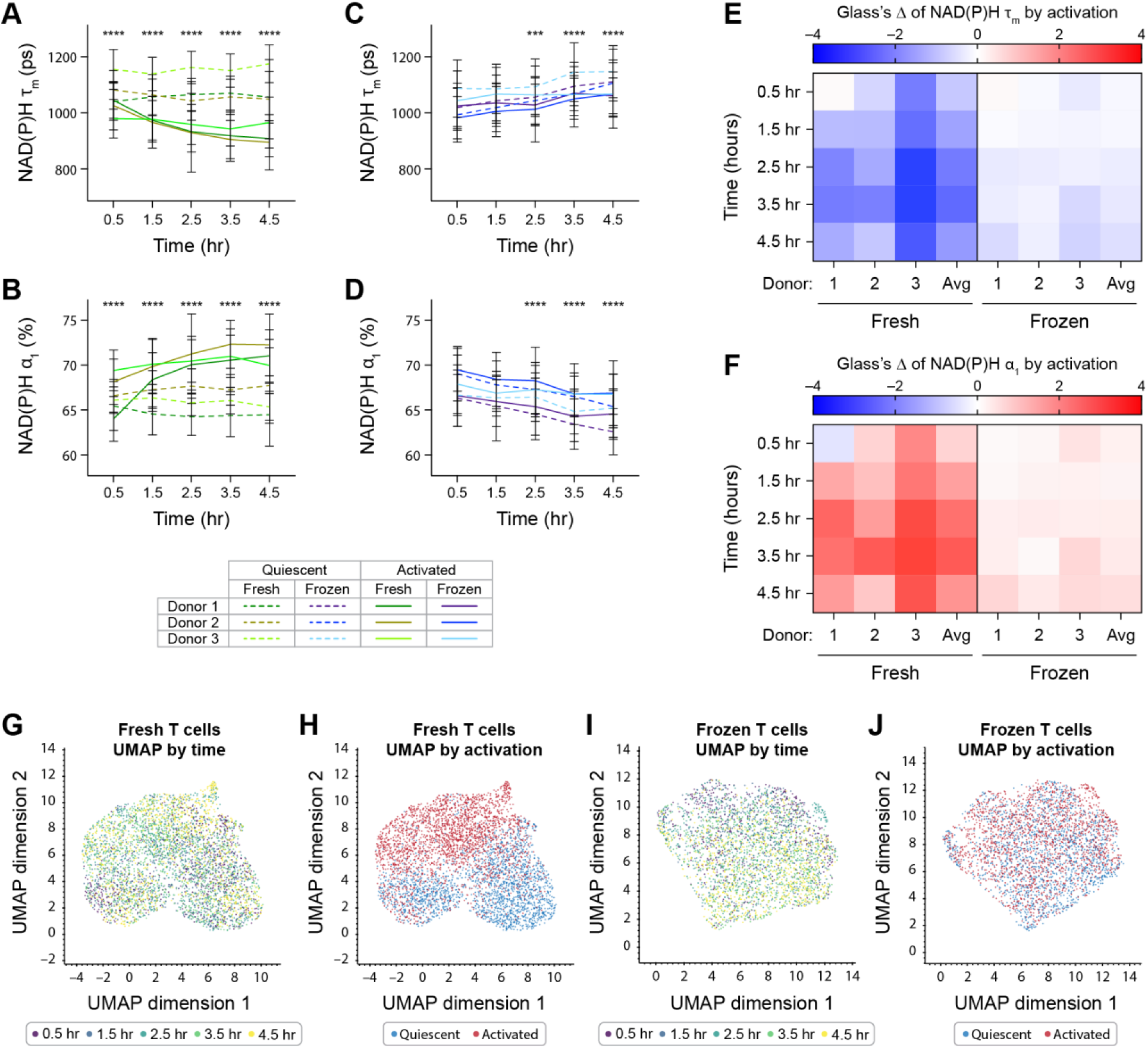
Cryopreservation delayed and diminished activation response in frozen T cells upon thawing. **(A-D)** Quantification of NAD(P)H τ_m_ and NAD(P)H α_1_ of **(A-B)** fresh or **(C-D)** frozen T cells from 3 independent donors. Lines represent donor averages, color coded by condition (fresh or frozen) and donor, while line patterns indicate activation status (dashed line: quiescent, solid line: activated). n = 296-618 cells/condition/timepoint across 3 donors. ANOVA with three factors: donor (donor 1, 2, and 3), time (0.5-4.5 hours), and activation status (quiescent, activated). Tukey post hoc test was used to determine statistical significance for multiple comparisons between quiescent and activated groups at each time point. **(E-F)** Glass’s Δs to quantify effect size of activation on **(E)** NAD(P)H τ_m_ and **(F)** NAD(P)H α_1_ of fresh and frozen T cells over time, with respect to corresponding quiescent group at each time point. **(G-H)** Uniform Manifold Approximation and Projection (UMAP) of 11 OMI parameters (NAD(P)H and FAD τ_m_, τ_1_, τ_2_, α_1_, α_2_, and cell size) of fresh T cells from 3 donors, color coded by **(G)** time and **(H)** activation status. n = 4595 cells. **(I-J)** UMAP of 11 OMI parameters of frozen T cells from 3 donors, color coded by **(I)** time and **(J)** activation status. n = 3451 cells. Bars are mean ± standard deviation. *** p < 0.001, **** p < 0.0001.

Compared to fresh T cells, donor-matched frozen T cells displayed smaller and delayed changes due to activation, with no significant differences in NAD(P)H τ_m_ or NAD(P)H α_1_ observed between frozen quiescent and frozen activated T cells until 2.5 hours post activation (**Fig 3C, D**). At the 2.5-hour time point, frozen activated T cells displayed significantly lower NAD(P)H τ_m_ and higher NAD(P)H α_1_ compared to frozen quiescent T cells. These differences continued up to 4.5-hours post activation (**Fig 3C, D**). Normalized NAD(P)H intensity of frozen T cells was less responsive to activating stimulus (**Supplementary Fig 2B**).

To determine the effect size of activation on fresh and frozen T cells, we calculated the Glass’s Δ at each time point with respect to donor-matched quiescent groups (**Fig 3E, F**). While we observed metabolic changes in a similar direction with activation in both fresh and frozen T cells, specifically lower NAD(P)H τ_m_ and higher NAD(P)H α_1_, the activation effects on fresh T cells (**Fig 3E, F – left**) were consistently greater than on donor-matched frozen T cells (**Fig 3E, F – right**). Activation induced strong effects on NAD(P)H τ_m_ and NAD(P)H α_1_ of fresh T cells, with effect sizes of up to -2.8 and 3.0 within 4.5 hours following activation. This indicated a difference of up to 3 standard deviations between donor-matched activated and quiescent cells. Meanwhile, for frozen T cells, activation effects on NAD(P)H τ_m_ and NAD(P)H α_1_ were smaller and remained within -0.7 to 0.7. Similarly, we observed greater effect size of activation on normalized NAD(P)H intensity of fresh T cells compared to frozen T cells (**Supplementary Fig 2C**). However, effect sizes due to activation did increase over time for frozen T cells, with the greatest Glass’s Δ observed at 4.5-hour post activation (**Fig 3E, F – right**). This suggests that as activation duration and time post-thaw increased, the ability of frozen T cells to respond to activating stimulus was recovered.

To understand the metabolic landscape of fresh and frozen T cells during activation, we projected all OMI lifetime parameters of fresh and frozen T cells with different activation statuses on a two-dimensional Uniform Manifold Approximation Projection (UMAP) (**Fig 3G-J**). We observed the OMI landscape of fresh T cells mainly clustered based on based on activation rather than by time (**Fig 3G-H, Supplementary Fig 2D, E**). Meanwhile, frozen T cells demonstrated a shift in clustering pattern from the top left to bottom right corner of the UMAP over the imaging time course, indicating a gradual shift in their metabolism (**Fig 3I),** but did not display distinct clusters based on activation status (**Fig 3J, Supplementary Fig 2F, G**). This suggests that within 4.5 hours upon stimulation, OMI captured more robust and significant metabolic changes in response to activation in fresh T cells compared to donor-matched frozen T cells. The distinct OMI metabolic profiles of fresh activated and quiescent T cells also allowed classification by activation status with high sensitivity and specificity (AUC > 0.91) across several classifier models (**Supplementary Fig 2H**), which could not be achieved in frozen T cells (AUC <0.68) (**Supplementary Fig 2I**). Additionally, we also observed significantly higher production of pro-inflammatory cytokine TNF-α by fresh activated T cells compared to donor-matched frozen activated T cells after 4.5 hours of stimulation (**Supplementary Fig 2J**). Overall, our data suggest a delayed and diminished activation response by frozen T cells compared to fresh T cells upon thawing.

### OMI identified activation in frozen T cells after 48 hours of stimulation

As frozen T cells showed delayed and diminished response to activating stimulus within the first 4.5 hours post-thaw, we further investigated the impact of cryopreservation and thawing on T cell activation response at a later time point. OMI at 48 hours post activation revealed significant metabolic and morphological differences between quiescent and activated T cells, for both fresh and frozen groups (**Fig 4A**). We observed significantly lower NAD(P)H τ_m_ (**Fig 4B, C**), higher NAD(P)H α_1_ (**Fig 4E, F**), greater cell size (**Supplementary Fig 3A, B**), as well as higher ORR (**Supplementary Fig 3D, E)** in fresh activated and frozen activated T cells compared to donor-matched quiescent cells. There was no significant difference in NAD(P)H τ_m_ between fresh and frozen cells at 48 hours; however, frozen activated T cells displayed slightly lower NAD(P)H α_1_ **(Fig 4D, G)**. Meanwhile, frozen quiescent T cells demonstrated greater cell size and lower ORR compared to fresh quiescent T cells (**Supplementary Fig 3C, F**). Overall, donor-matched Glass’s Δ revealed consistently great activation effects (|Glass’s Δ| >3) at 48 hours in both fresh and frozen groups (**Fig 4B, C, E, and F**). These findings suggested that at 48 hours after thawing and activation, frozen T cells recovered their activation response. However, we did observe consistently lower fold-expansion in frozen T cells compared to their donor-matched fresh counterparts throughout a 7-day expansion period (**Fig 4H**), which indicated a potential impact of cryopreservation on T cell expansion capacity.

**Figure 4.**
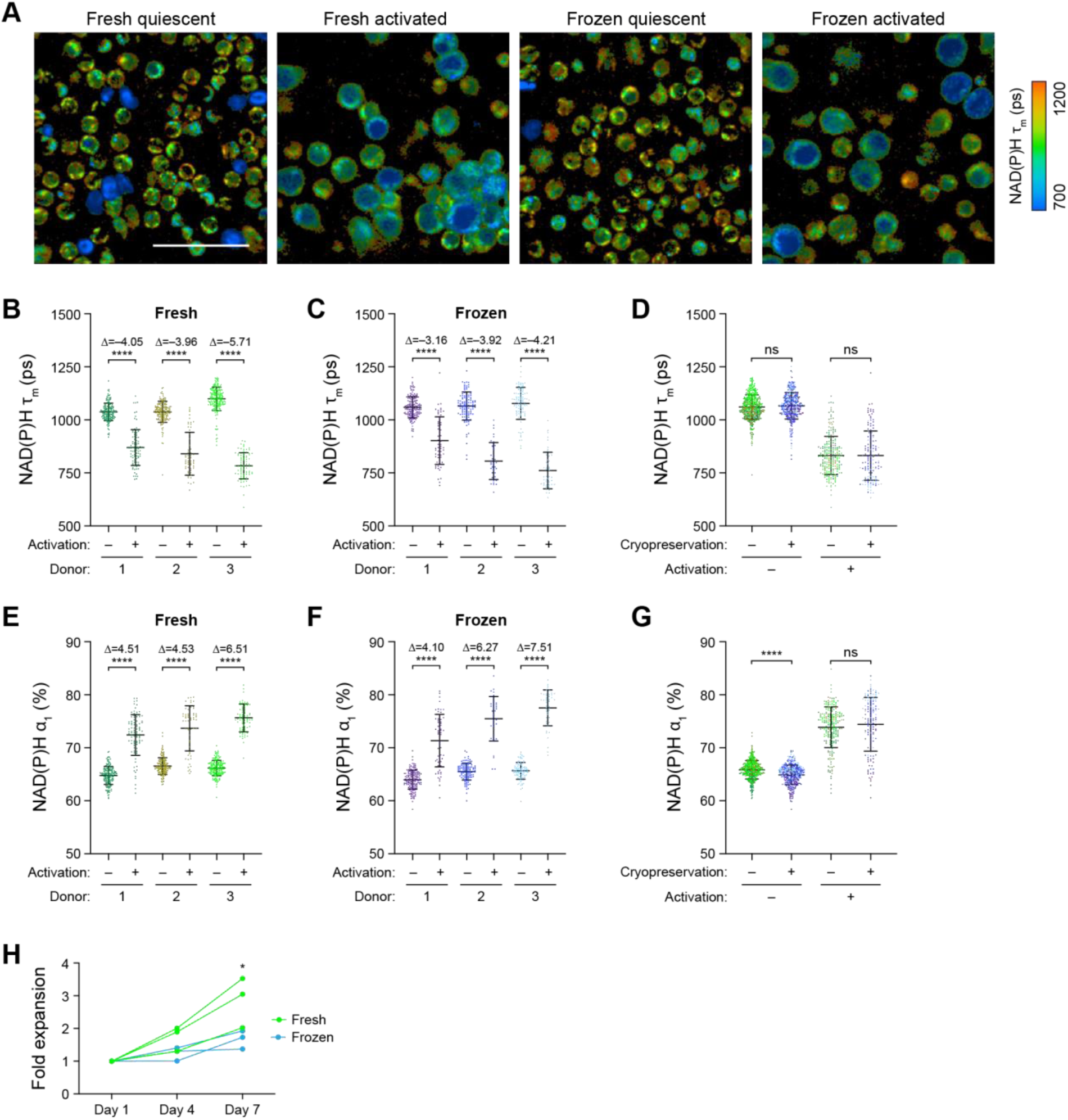
Activation response of frozen T cells recovered after 48 hours of activation post-thaw. **(A)** Representative NAD(P)H τ_m_ images of fresh and frozen quiescent and activated T cells at 48 hours after activation. **(B-C)** Quantification and **(D)** comparison of NAD(P)H τ_m_ of **(B)** fresh and **(C)** frozen quiescent and activated T cells at the 48-hour time point from 3 donors. **(E-F)** Quantification and **(G)** comparison of NAD(P)H α_1_ of **(E)** fresh and **(F)** frozen quiescent and activated T cells at the 48-hour time point. For **(B-G)** n = 36-195 cells/condition/donor. Two-sided non-parametric Kruskal-Wallis test with Dunn’s post hoc tests for multiple comparisons between **(B, C, E and F)** quiescent versus activated groups for each donor and **(D, G)** fresh versus frozen for all 3 donors. **(H)** Fold expansion through a 7-day expansion time course of fresh and frozen activated T cells from 3 donors. Two-way repeated measures ANOVA with two factors: day (day 1, 4, and 7) and cryopreservation status (fresh, frozen). Sidak post hoc test for multiple comparisons between fold expansions of fresh versus frozen T cells at each time point. 3 data points from 3 independent donors were counted as repeated measurements for statistical test. Scale bar is 50μm. Bars are mean ± standard deviation. * p < 0.05, **** p < 0.0001.

Binding of stimulatory and co-stimulatory receptors on T cells, such as via CD3/CD28 antibody, provides activation signals for T cells. Meanwhile, antigen-specific activation serves as another mode of activation that is relevant for *in vivo* functions of adoptive cell therapies including CAR and TCR therapies. Therefore, we also further investigated antigen-specific activation response in frozen T cells since cryopreservation is widely used in the context of these adoptive cell therapies. We performed OMI of CMV-specific T cells activated with StemCell αCD2/αCD3/αCD28 or HLA-matched CMV peptides. We observed similar metabolic response in frozen CMV-specific T cells upon thawing, characterized by increased in NAD(P)H τ_m_ that was sustained up to 5 hours post-thaw (**Supplementary Fig 4A-D**). This is consistent with our data on frozen CD3 T cells reported above. Similarly, over the first 5 hours of activation post-thaw, CMV-specific T cells activated with αCD2/αCD3/αCD28 antibody and CMV-peptide both showed significant increase in NAD(P)H τ_m_ (**Supplementary Fig 4E, F**). This suggests a potential impact of cryopreservation and thawing not only on antibody-activation but also antigen-specific activation response in T cells. Notably, while low NAD(P)H τ_m_ is characteristic of T cell activation response, CMV-specific T cells activated with CMV peptide displayed slightly higher NAD(P)H τ_m_ compared to the quiescent group during the first hour post-thaw (**Supplementary Fig 4G**). Interestingly, compared to αCD2/αCD3/αCD28 antibody activation, antigen specific activation with HLA-matched CMV-peptide induced faster and stronger metabolic response in frozen CMV-specific T cells (**Supplementary Fig 4G -pink line versus purple line, Supplementary Fig 4H – right versus left**). Starting at 3 hours post-thaw, frozen CMV-specific T cells activated with HLA-matched CMV peptide showed significantly lower NAD(P)H τ_m_ compared to control quiescent group. This difference was sustained up to the 5-hour time point, and Glass’s Δs representing antigen-specific activation effects on NAD(P)H τ_m_ strengthened overtime (Δ = -0.03 at 3 hours post-thaw, Δ = -0.44 at 5 hours post-thaw). Meanwhile, frozen CMV-specific T cells activated with αCD2/αCD3/αCD28 antibody did not show significantly lower NAD(P)H τ_m_ compared to control group until the 5-hour time point (**Supplementary Fig 4G – purple line, Supplementary Fig 4H – left**). As T cells produce several inflammatory cytokines such as IFN-γ upon activation, we further quantified the amount of IFN-γ secreted by different CMV-specific T cells groups during the 5-hour activation time course. Interestingly, antigen specific activation with CMV peptides resulted in significantly higher IFN-γ production compared to antibody activated cells and quiescent cells (**Supplementary Fig 4I**). This is consistent with the greater changes in NAD(P)H τ_m_ observed in the antigen-activated group. In summary, our data suggest that cryopreservation affects both CD3 receptor activation and antigen-specific activation in T cells. However, antigen-specific activation induces stronger and faster metabolic as well as functional response in post-thaw T cells.

### OMI characterized early metabolic stress in cryopreserved cancer patient T cells post-thaw

Cryopreservation is essential for the storage and distribution of cell therapies. Using OMI, we have shown that short-term cryopreservation (24 hours) significantly alters the metabolic state of T cells from healthy donors during the early post-thaw period. Therefore, we aimed to further assess the impacts of cryopreservation on T cells from four patients with either diffuse large B cell lymphoma (DLBCL) or mantle cell lymphoma (MCL), who would later undergo bispecific CD20/CD19 CAR T cell therapy (**Supplementary Table 1**)^35^. A subset of CD3 T cells isolated from these patients as the starting materials for their CAR T therapy were cryopreserved in Cryostor_media containing 5% DMSO at -80^0^C. Using OMI, we monitored metabolic profiles of these patient T cells immediately after thawing and over the course of 4.5 hours post-thaw. We identified distinct metabolic responses in cryopreserved T cells from healthy donors (**Fig. 5A,B**) versus lymphoma patients (**Fig. 5C,D**). Notably, a second population emerged over time in patient T cells -characterized by a low ORR, increased NAD(P)H α_1_ and reduced NAD(P)H τ_m_ -which was not observed in healthy donor T cells (**Fig 5B, D**). This population also exhibited morphological signs of stress, including compromised nuclear and cellular membrane integrity (white arrow, **Fig 5C**).

**Figure 5.**
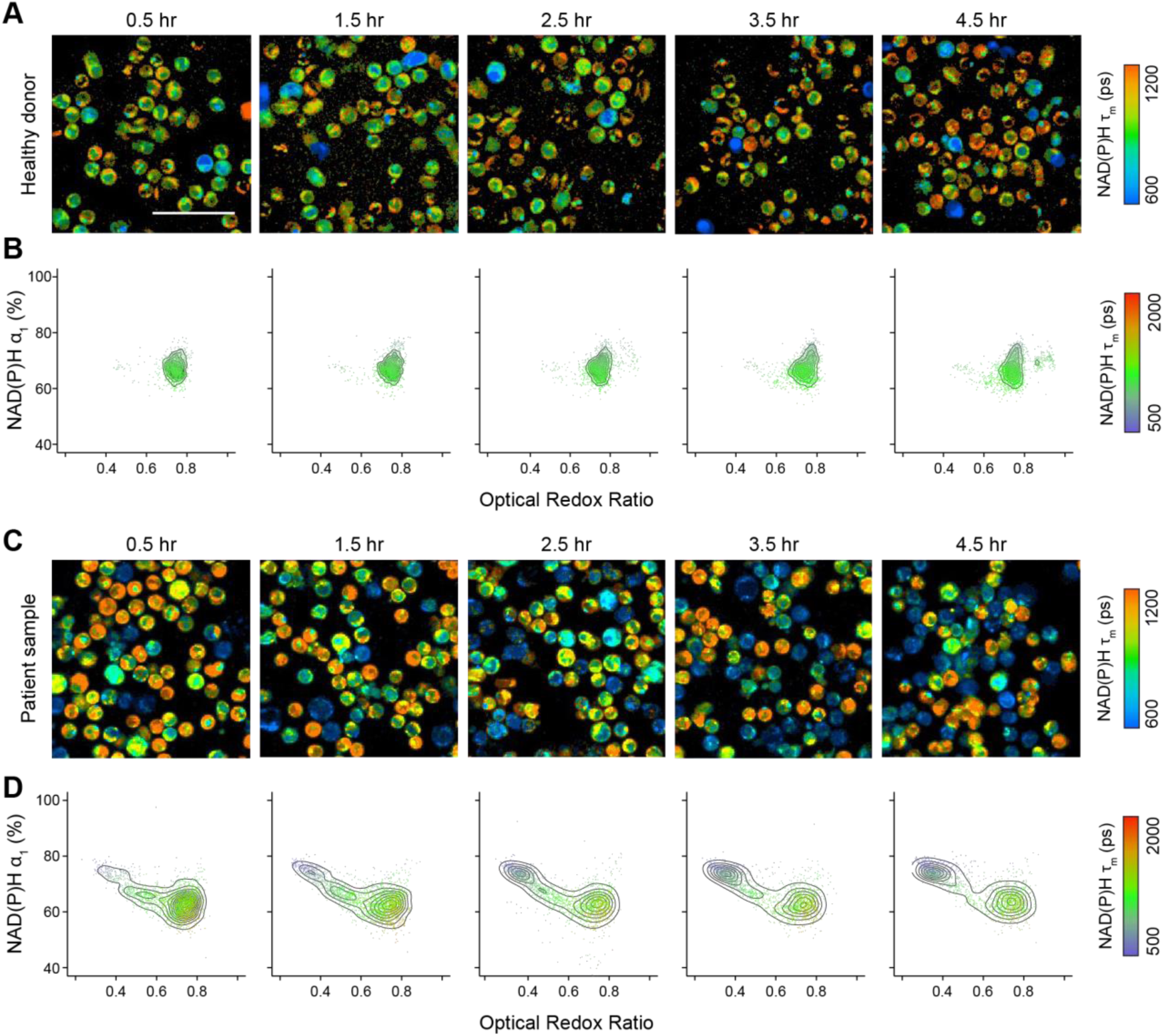
Cryopreserved CD3 T cells from healthy donors and cancer patients exhibited distinct metabolic responses upon thawing. **(A)** Representative NAD(P)H τ_m_ images and **(B)** Two-dimension density contour of NAD(P)H α_1_ and ORR, color coded by NAD(P)H τ_m_ of CD3 T cells from 3 healthy donors throughout 4.5 hours upon thawing. **(C)** Representative NAD(P)H τ_m_ images and **(D)** Two-dimension density contour of NAD(P)H α_1_ and ORR, color coded by NAD(P)H τ_m_ of CD3 T cells from 4 patients with either diffuse large B cell lymphoma or mantle cell lymphoma throughout 4.5 hours upon thawing. n = 8046 cells for **(B)** and 11921 cells for **(D)**. Scale bar is 50μm.

We hypothesize that this population reflects an early metabolic stress response induced by cryopreservation and thawing. Density distribution of NAD(P)H α_1_ and ORR revealed the progressive transition from low-NAD(P)H α_1_ and high-ORR to high-NAD(P)H α_1_ and low-ORR in these patient T cells over 4.5 hours upon thawing (**Fig 6A, B**). To identify these metabolic subpopulations, we applied a two-component Gaussian mixture model to the density distributions of NAD(P)H α_1_ and ORR pooled from four patients and defined the thresholds for each subpopulation as 68.96 for NAD(P)H α_1_ and 0.61 for ORR (**Fig 6C, D**). Cells within this gate (NAD(P)H α_1_ > 68.96% and ORR < 0.61) were considered metabolically stressed, while those outside of this gate represented a metabolically fit population upon thawing (**Fig 6E**). Using this OMI-based gating, we observed a time-dependent decline in metabolic fitness within the first 4.5 hours post-thaw from patient T cells (**Fig 6F**). Remarkably, metabolic fitness assessed by OMI at 4.5 hours were consistent with viability measured by Trypan blue exclusion at 24 hours in patient samples (**Fig. 6G**). Hence, in this context, we defined “metabolic fitness” as the ability to recover and retain viability at 24 hours post-thaw. This suggests that OMI could serve as an early, non-destructive indicator of T cell recovery potential post-thaw.

**Figure 6.**
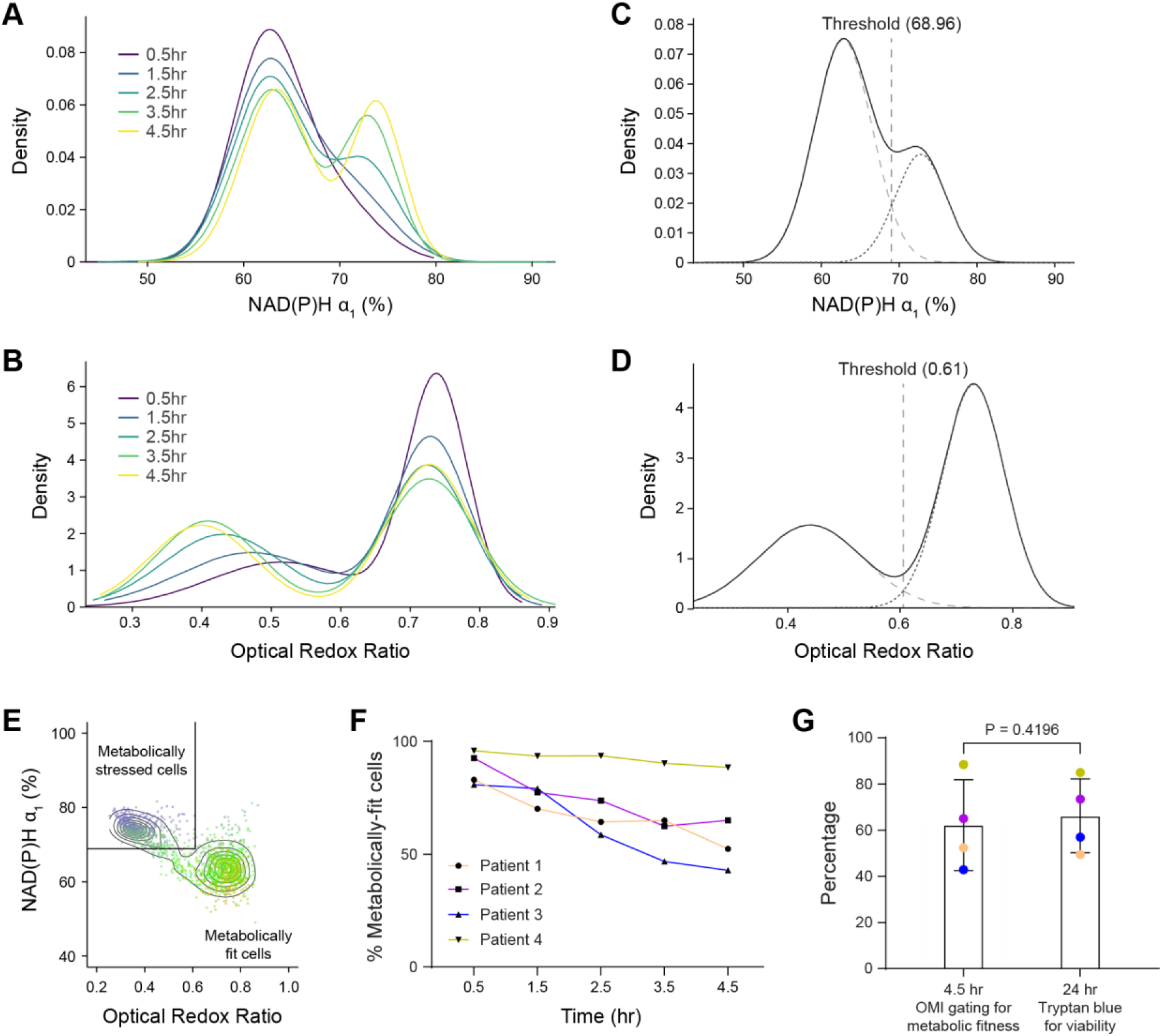
OMI provides an early measurement of recovery potential post-thaw in patient T cells. (**A-B**) Density distribution of (**A**) NAD(P)H α_1_ and (**B**) ORR in patient T cells over a 4.5-hour post-thaw time course. (**C-D**) Two-component Gaussian mixture models fit to pooled data across all time points from four patients for (**C**) NAD(P)H α_1_ and (**D**) ORR. Thresholds were determined as the intersection of the two fitted Gaussian distributions. n = 11921 cells **(E)** Gating strategy applied to NAD(P)H α_1_ and ORR to define metabolically stressed and fit cells. (**F**) Percentage of metabolically-fit cells gated by OMI features (NAD(P)H α_1_ and ORR 4.5 hours post thaw with cell viability by Trypan blue staining at 24 hours. n = 4 patients. Two-sided paired t-test.

### OMI identified activation-associated metabolic changes in T cells from complete responders post-thaw

Prior studies have shown that the metabolism and function of patient T cells used as starting materials can influence clinical outcomes following CAR T cell therapy^36^. Therefore, based on previously defined OMI features, we gated for metabolically fit T cells from starting materials of four patients and evaluated their response upon stimulation with αCD2/αCD3/αCD28 during the early post-thaw period. Interestingly, metabolically fit T cells from patients who later achieved complete response (CR) to CD20/CD19 CAR T cell treatment demonstrated activation signatures over the 4.5-hour time course. Specifically, activated T cells from these patients showed significantly higher NAD(P)H α_1_ compared to donor-matched quiescent cells starting at 2.5 hours post-thaw (**Fig. 7A**), while no significant activation-associated changes in NAD(P)H α_1_ were observed in activated T cells from patients with later partial response (PR) or progressive disease (PD) (**Fig 7B,C**). Additionally, activated T cells from CR patients also exhibited decreased NAD(P)H τ_m_, increased ORR, and increased cell size compared to quiescent cells throughout the 4.5hour post-thaw time course (**Fig 7D-F**). Previous studies have validated the association between these OMI features and activation response in T cells, suggesting that metabolically fit cells from CR patients retained the ability to activate post-thaw^31,32,37^. Meanwhile, T cells from PR and PD patients did not show these clear activation characteristics with no significant difference observed in activated versus quiescent groups, suggesting while these cells retained viability following the freeze-thaw process, they demonstrate impaired functions (**Fig 7C-F**).

**Figure 7.**
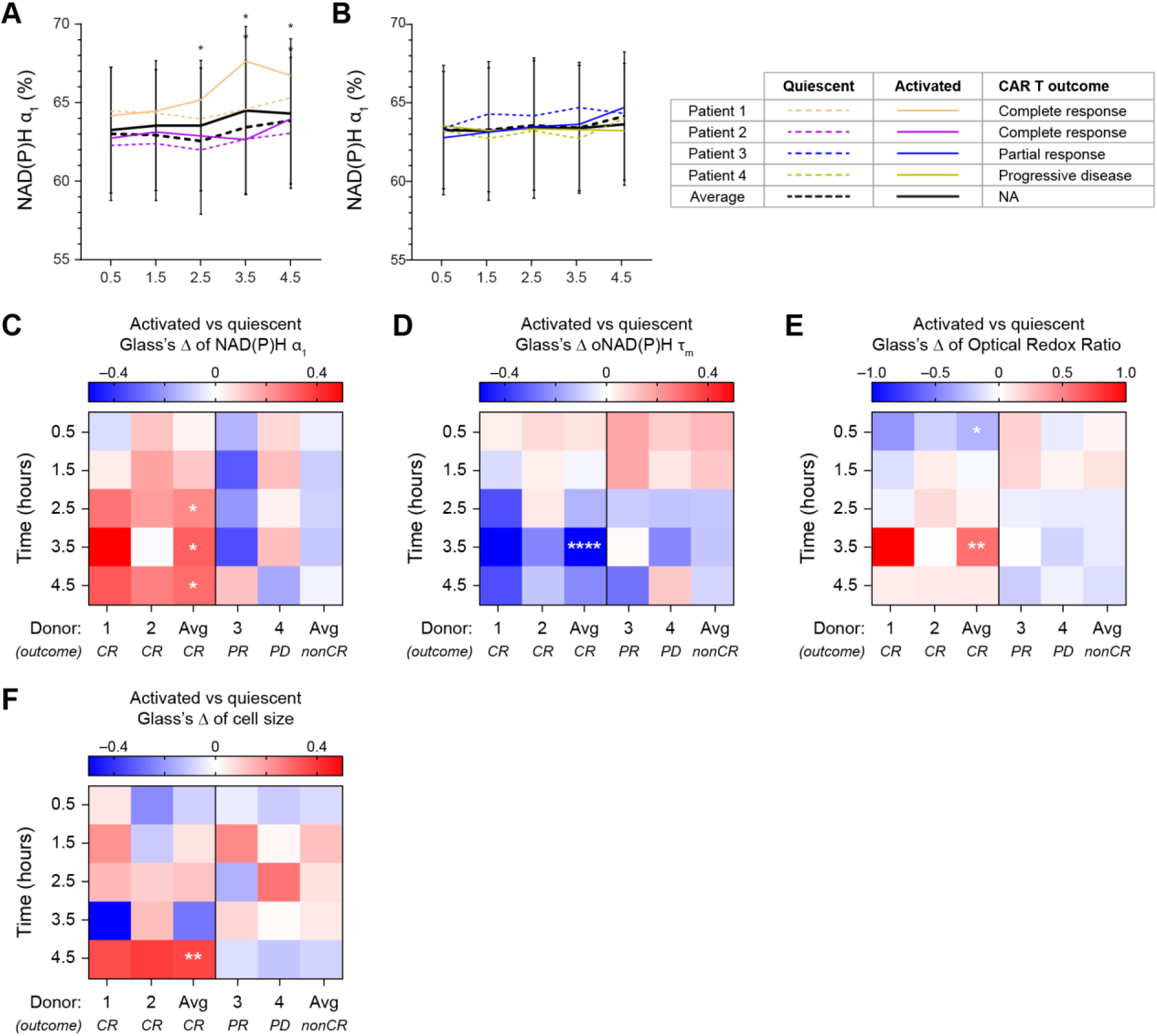
Metabolically fit T cells from patients achieving complete response to CAR T treatment demonstrated activation response post-thaw. **(A-B)** NAD(P)H α_1_ of quiescent and activated T cells from starting materials of patients with **(A)** complete response (CR) or **(B)** partial response (PR) and progressive disease (PD) in response to later CD20/CD19 CAR T treatment. Lines represent donor average, color coded by donors, patterned by activation status (dashed line: quiescent, solid line: activated). n = 197-367 quiescent cells and 157-290 activated cells at each time point across 2 donors (patient 1 and patient 2) for **(A)**, n = 257-576 quiescent cells and 314-571 activated cells at each time point across 2 donors (patient 3 and patient 4) for **(B)**. **(C-F)** Glass’s Δs to quantify effect size of activation on **(C)** NAD(P)H α_1_**, (D)** NAD(P)H τ_m_, **(E)** ORR, and **(F)** cell size of metabolically fit T cells from 4 patients with respect to donor-matched quiescent group at each time point. *: p < 0.05. **: p < 0.01, ****: p < 0.0001. Linear model with activation status as fixed effect and donor as blocking factor with Benjamin-Hochberg post-hoc test for multiple comparisons between activated and quiescent groups at different timepoints. For (A-B), error bars are standard deviation across 2 CR and 2 non-CR patients.

## DISCUSSION

Cryopreservation of biological samples is central in research and clinical translation of cell therapies, where the intricate manufacturing process requires transportation of cell products between the manufacturing facility and treatment site. However, cryopreservation that retains cell viability and function remains challenging for several primary immune cells such as neutrophils and natural killer cells, limiting their clinical translation. Previously, cryopreservation has been shown to impair metabolism and reduce immunomodulatory functions of mesenchymal stem cells due to heat-shock response, which potentially compromises their therapeutic benefits^38^. Meanwhile, resting of cryopreserved natural killer cells overnight post-thaw improves their cytotoxic functions^39^, suggesting that the period immediately post-thaw is a critical time window when cell function might be sensitive to cryoinjury. Since the activation of cryopreserved cells upon thawing is involved in several stages of CAR T cell manufacturing and treatment, we used OMI, a label-free non-invasive imaging method, to characterize the impact of cryopreservation and thawing on T cell activation response throughout the first 4.5 hours post-thaw. Across three healthy donors, we showed a conserved metabolic response of T cells upon thawing with high NAD(P)H τ_m_ and low NAD(P)H α_1_ that was sustained up to 4.5 hours. These shifts in OMI measurements indicate an increase in NAD(P)H binding activity, potentially due to cell metabolism ramping up after thawing^40^. Meanwhile, fresh quiescent T cells exhibited stable OMI measurements throughout the imaging time course. We hypothesized that T cell metabolism immediately upon thawing was not ready to support proper activation response. Our findings confirm a delayed, diminished activation response in frozen T cells immediately post-thaw, further supporting our hypothesis. Additionally, our data are consistent with previous studies showing impaired immune response, specifically reduced cytokine production, due to cryopreservation in peripheral blood mononuclear cells^41^. Interestingly, while activation response in frozen T cells was recovered at 48 hours post-thaw, their expansion capacity throughout a 7-day expansion process remained lower than donor-matched fresh T cells. This suggests that impairment in the early activation response could negatively impact T cell expansion.

Using frozen CMV-specific T cells and HLA-matched CMV-peptides, we demonstrated the impact of cryopreservation on both CD3-mediated and antigen-specific activations in T cells. We did observe faster and stronger metabolic changes, along with increased production of inflammatory cytokines in frozen CMV-specific T cells activated with CMV-peptide compared to CD3-antibody. Our findings indicate that OMI is sensitive to differences in T cell metabolic response towards two types of activating stimuli.

Many CAR T manufacturing processes currently rely on cryopreservation at several stages, with some protocol demanding freezing of the starting material before shipping to the manufacturing site; however, the impacts of cryopreservation on patient T cells prior to CAR T manufacturing have yet to be fully characterized. Additionally, while previous research on the impact of cryopreservation on the efficacy of CAR T cell therapy suggests that cryopreserved products offer comparable potency and treatment response, there have been some evidence of lower CAR T persistence in patients receiving frozen products, suggesting that this effect could be donor-dependent or CAR-T model dependent^42^. Cryopreservation has also been shown to impair mitochondrial coupling efficiency and ATP production in leukocytes, while increasing dependence on glycolysis^43^. However, these effects were characterized after overnight resting in leukocytes. Using OMI, we identified early metabolic stress signals induced by the freeze-thaw process in patient T cells and developed a label-free gating strategy for metabolic fitness, which is defined as the ability to recover and the maintain viability at 24 hours post thaw. OMI analysis of metabolically fit cells upon stimulation with activating antibodies reveals functional differences in T cells isolated from patients achieving CR compared to those obtained from PR/PD patients, although the sample size (n=4 patients) is too low to claim the generalizability of these findings. Activation signatures, including high NAD(P)H α_1_, low NAD(P)H τ_m_, low ORR, and increased cell size, were observed in CR patient T cells while absent in those from PR/PD patients. This suggests that retention of viability might not be sufficient for functional response in T cells post-thaw, and that metabolic features should be assessed to evaluate patient T cell quality for CAR T manufacturing. Additionally, these findings support the application of OMI as a label-free and single-cell analytical tool for continuous monitoring of the dynamic and heterogenous response in patient T cells. As these cells were undergoing both metabolic stress due to freeze-thaw process and activation response to stimulating antibodies, OMI allowed functional characterization of each subpopulation that could otherwise be confounded in bulk population measurements. Our data highlight the potential of OMI as a sensitive tool for characterizing patient T cell quality after cryopreservation and screening of patient T cell fitness for CAR T manufacturing.

There are limitations to this study. First, it is important to note that several factors potentially contribute to the differences in metabolic responses of T cells from healthy donors and lymphoma patients upon thawing, including the patient disease status and previous lines of treatments, as well as different T cell isolation and cryopreservation protocol. Therefore, while OMI was able to identify distinct metabolic signatures from these samples, further research with controlled experimental conditions across more healthy donors, a larger patient cohort, and additional CAR T models are needed to validate our findings and to evaluate how these factors impact T cell function and recovery potential post cryopreservation. Additionally, different cryopreservation and thaw techniques are used in different contexts of T cell manufacturing, and this study did not exhaustively test or compare these methods. In order to determine the optimal cryopreservation methods and post-thaw condition for specific cell therapy applications, OMI measurements should also be benchmarked with other metabolic and functional assays to better characterize the complex response of T cells post-thaw. Our findings also focused on cryopreservation of peripheral T cells prior to CAR manufacturing, while cryopreserved final products for storage and distribution is a more common practice in the field. Hence, future studies will focus on characterizing frozen CAR T products to better understand its impact on their therapeutic efficacy and clinical outcomes. Finally, cryopreservation is also critical for scientific research and other medical applications besides cell therapy (such as in vitro fertilization or testing of potential pathogens)^44^. While we did observe significant antigen response from CMV-specific T cells upon thawing, future research will further investigate whether OMI is sensitive to the impacts of cryopreservation in these contexts.

## MATERIALS AND METHODS

### T cell isolation and culture

Healthy donors were recruited and informed consent was collected from all donors under a protocol approved by the Institutional Review Board at the University of Wisconsin – Madison. CD3 T cells were isolated from peripheral blood of healthy donors using RossetteSep Human T cell enrichment cocktail (STEMCELL Technologies) following the manufacturer’s protocol. Briefly, peripheral blood was incubated with 50μL/mL T cell enrichment cocktail, then diluted in PBS + 2% Fetal Bovine Serum (FBS) and layered on Lymphoprep density gradient (STEMCELL Technologies) for separation using centrifugation. Isolated T cells were then divided into two groups for fresh culture or cryopreservation. For the fresh culture group, T cells were resuspended in ImmunoCult XF T cell expansion medium at 1 million cells/mL and cultured overnight at 37C, 5% CO2. For cryopreservation, T cells were resuspended in FBS+10% DMSO at 1 million cells/mL. Cryopreservation vials containing T cells were then encapsulated in a freezing container (CoolCell LX Cell Freezing Container, Corning) to control for freezing rate (-1C/minute) and transferred to -80C freezer to be frozen overnight (**Supplementary Fig 1**).

### Isolation and cryopreservation of patient T cells

Patient cells were collected by clinical leukapheresis; CD4/CD8 selection was performed on a CliniMACS Prodigy instrument (Miltenyi Biotec, Bergisch Gladbach, Germany) by positive immunomagnetic separation. 100-150E6 CD4/CD8-positive cells were washed with PBS, and then resuspended in 4 mL Cryostor CS5 (Biolife Solutions, Bothell, WA). The resuspended cells were placed in a 4 mL cryovial and frozen at -80C in a Mr. Frosty device (Thermo Scientific, Waltham, MA).

### Thawing and activation of T cells

Cryopreservation vials were submerged in a water bath at 37C for 30 seconds. For healthy donor T cells, warm fresh ImmunoCult XF T cell expansion medium was added to the cryopreservation vial one drop at a time. When no ice crystals were visible, all T cells were then transferred into 10mL of warm fresh ImmunoCult XF media, then centrifuged to wash out DMSO residue. T cells were then counted and plated at 200,000 cells in 200 μL of fresh media into a glass-bottom 96 well plate. Immediately upon thawing and plating, frozen T cells were activated with 5μL of StemCell αCD2/αCD3/αCD28 antibody (**Supplementary Fig 1**). To maintain consistency with our bispecific CD20/CD19 CAR T manufacturing condition, T cells from patients with B cell malignancies were thawed following the same procedure using fresh warm TexMACs supplemented with 3% human serum and 200U/mL IL-2 to wash out residual DMSO and activated with TransAct αCD3/αCD28 antibody.

For the antigen specific experiment, HLA-A*0201 restricted anti CMV T cells were obtained from Celero (item #1049-5147JN21and item #1049-5085AP21). Upon arrival, T cells were kept in liquid nitrogen until used. T cell thawing was performed following a similar protocol as above. Briefly, RPMI + 2% FBS + 1% Pen/Strep was warmed up to 37C, then added by drop into cryopreserved T cell vials. T cells were then washed once with fresh warm media and plated at a density of 1 million cells/mL. Immediately upon thawing, CMV-specific T cells were plated onto 35mm glass bottom, Poly-D-lysine coated imaging dishes (MatTek) at a concentration of 200,000 cells/200 μL in Immunocult T cell expansion media. T cells were then stimulated with either 5μL of StemCell αCD2/αCD3/αCD28 antibody or 1μL iTAg Tetramer/APC–HLA-A*02:01 CMV pp65 peptide (NLVPMVATV) (MBL International) to generate antibody and CMV activated groups, respectively.

### T cell imaging with OMI

30 minutes after thawing and activation, T cells were imaged using OMI every hour up to 4.5 hours in a stage top incubator (37°C, 5% CO_2_) (**Supplementary Fig 1**) as previously described^37^. OMI was performed on a multiphoton microscope (Ultima, Bruker) using an inverted laser-scanning microscope body (Ti-E, Nikon) equipped with an ultrafast tunable laser source (Insight DS+, Spectsra Physics). Two-photon excitation of NAD(P)H and FAD were performed at 750nm (2.5 mW) and 890nm (4.5mW), respectively, using a 40X water immersion 1.15 NA objective (Nikon) with 2.5x optical zoom. Other imaging parameters include 4.8µs pixel dwell time, 60s integration time, and image size of 256 x 256 pixels. NAD(P)H and FAD emission spectra were collected using GaAsP photomultiplier tubes (H7422, Hamamatsu) using a 440/80 nm and 550/100nm bandpass filters, respectively. Fluorescence decays of NAD(P)H and FAD were acquired using time-correlated single-photon counting (TCSPC) electronics (SPC 150, Becker & Hickl GmbH) using Prairie View Software (Bruker). APC-conjugated CMV peptide was excited at 980nm and emission signal was collected using 690/50nm filter. Fluorescence intensity and lifetime images of NAD(P)H and FAD, together with immunofluorescence images of APC-conjugated CMV peptide, were collected for each field of view (FOV), with 3–5 representative FOVs imaged per condition. The instrument response function was measured using second-harmonic generation signal from urea crystals excited at 890nm.

### Image segmentation and OMI analysis

Fluorescence decay curves collected with TCSPC were fit to a double-exponential decay using the weighted-least square model in SPCImage software to determine pixel-wise lifetimes of free and protein-bound NAD(P)H and FAD. NAD(P)H and FAD lifetime images were binned to combine fluorescence counts from 25 neighbor pixels for fitting. Pixel-wise ORR was calculated as the normalized ratio between NAD(P)H fluorescence intensity and the sum of NAD(P)H and FAD fluorescence intensity. Individual cell and individual cytoplasm were segmented for CD3-mediated and antigen-specific experiments, respectively. CellPose cyto2 model was used for automatic segmentation of whole cells, with manual check by users to ensure accurate masking for each FOV. 13 OMI variables were collected and quantified, including: ORR, fluorescence intensity and mean fluorescence lifetime (τ_m_) of NAD(P)H and FAD, their free- and protein-bound lifetime components (τ_1_, τ_2_) and corresponding fractional contributions (α_1_, α_2_).

### Statistical analysis

Statistical significance among experimental groups was performed in Prism (v10.2.2) or R-Studio (v3.5.3). Two-sided non-parametric Kruskal-Wallis test was used to eliminate bias and assumption of normality in the data set. Dunn’s post hoc test was chosen to adjust for multiple comparisons. To determine the effect size of metabolic changes within the same treatment condition (e.g., frozen quiescent, frozen activated, fresh quiescent, or fresh activated group), Glass’s Δ was calculated as: 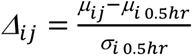 where *μ*_𝑖𝑗_ is the mean value for the OMI parameter of interest for donor *i* at time point *j*, *µ*_𝑖 0.5ℎ𝑟_ is the donor-matched mean value of the corresponding OMI parameter at the 0.5 hour time point, and *σ*_𝑖 0.5ℎ𝑟_ is the standard deviation of *μ*_𝑖 0.5ℎ𝑟_. To determine the effect size of activation on T cell metabolism at each time point, 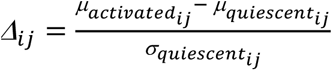. Glass’s Δ values were reported for individual donors, as well as the averaged values across three donors. A two-component Gaussian mixture model was fit to the density distribution of OMI parameters, such that each component has at least 10% weight of the total population. Thresholds were determined as the intersection of the two component densities using Brent’s method for root finding routine^45^.

## Data availability

All data are available upon request to the authors.

## Acknowledgement

We thank patients who enrolled in the study and the Cell Therapy Shared Resource at the Medical College of Wisconsin Cancer Center for the patient samples. Thank you to Matthew Stefley for graphic edits. Thank you to the Skala lab members for helpful discussion and comments on the manuscript.

## Author contributions

Conceptualization: D.L.P and M.C.S; Investigation and Methodology: D.L.P, M.K., C.W., and A.H.; Formal analysis and Software: D.L.P, W.Z.; Resources: A.G., T.K., P.H, and N.S.; Supervision: M.C.S; Writing-original draft: D.L.P and M.C.S; Writing-review & editing: D.L.P, M.K., A.H., A.G., W.Z, T.K., P.H, N.S.; and M.C.S.

---

**Supplementary Figure 1.**
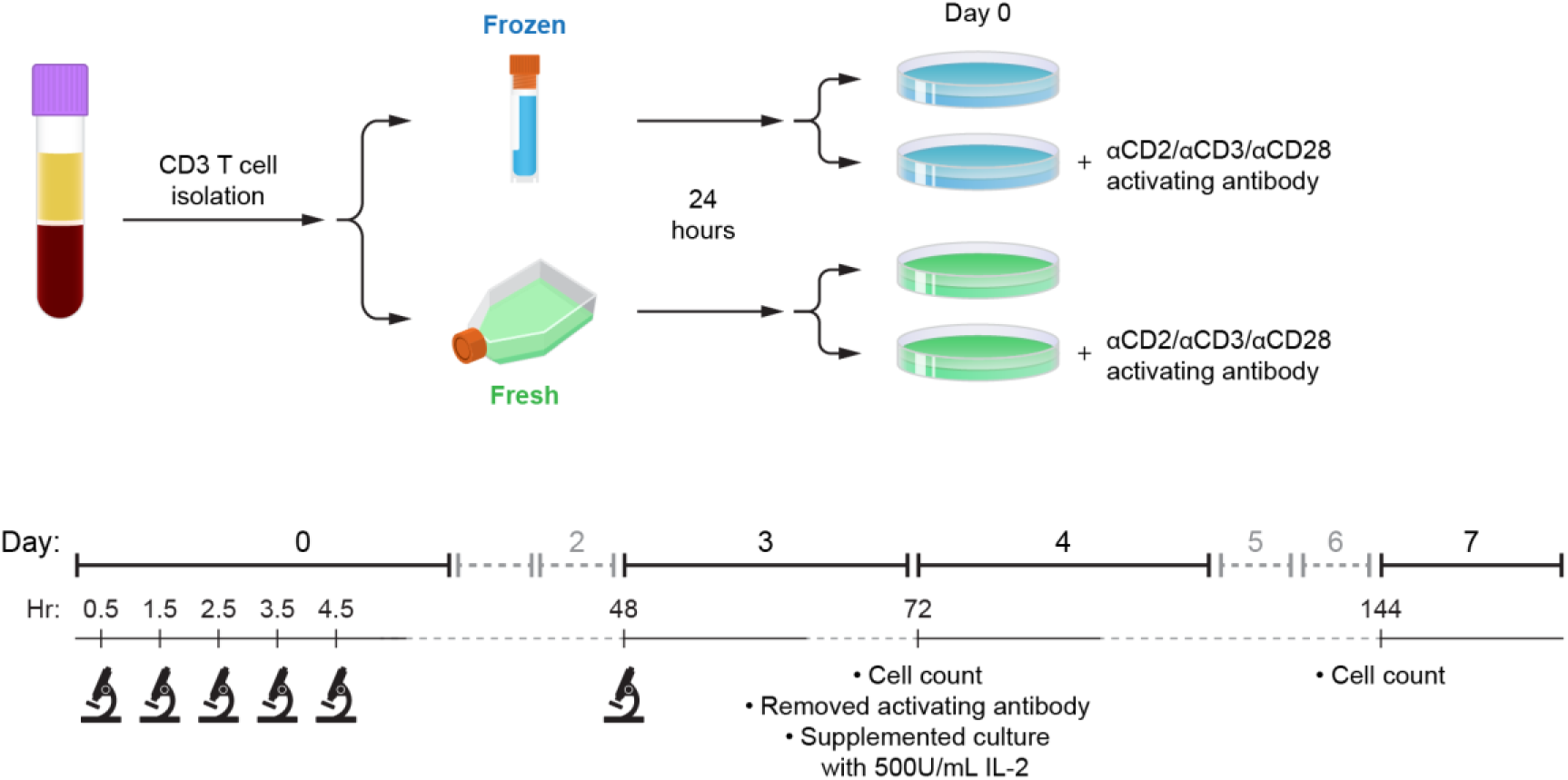
Experimental Setup. On day 0, CD3 T cells were isolated from peripheral blood of healthy donors and divided into two groups for cryopreservation or fresh culture overnight. After 24 hours (day 1), cryopreserved T cells were thawed, and donor-matched fresh T cells were also harvested. Both frozen and fresh T cells were stimulated with aCD2/aCD3/aCD28 T cell activator and imaged with OMI every hour up to 4.5 hours. Fresh and frozen quiescent and activated cells were imaged again at 48 hours (day 3). 72 hours post activation (day 4), cells were counted and resuspended in fresh ImmunoCult XF T cell expansion media supplemented with 500U/mL IL-2. Cells were expanded up to day 7, when cell count was performed to determine fold expansion. Created with BioRender.com.

**Supplementary Figure 2.**
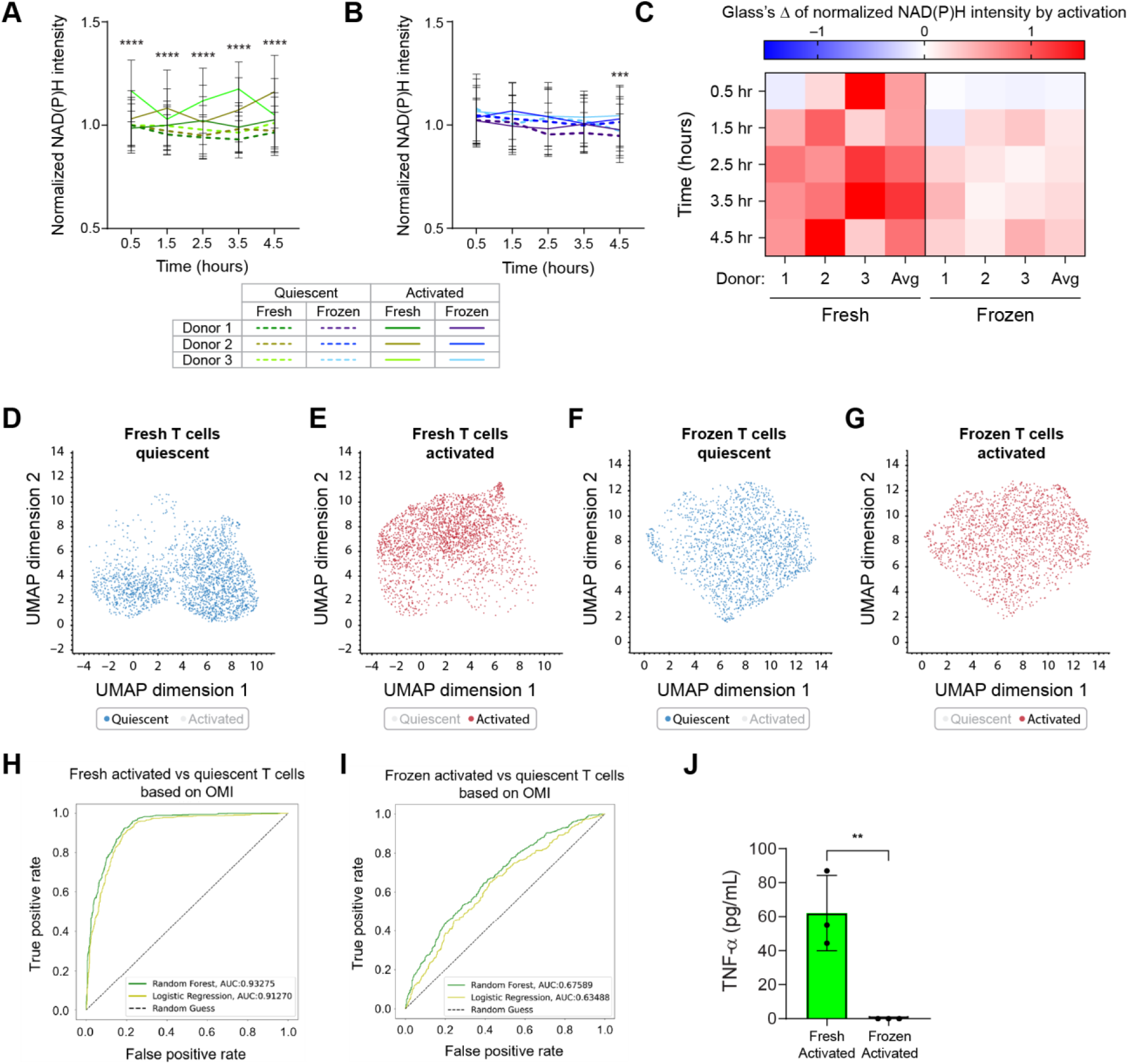
Frozen T cells demonstrated diminished activation response compared to donor-matched fresh T cells throughout 4.5-hour imaging time course. **(A-B)** Quantification of normalized NAD(P)H intensity of **(A)** fresh or **(B)** frozen quiescent and activated T cells from 3 independent donors. Lines represent donor averages, color coded by condition (fresh or frozen) and donor, line patterns indicate activation status (dashed line: quiescent, solid line: activated). NAD(P)H intensity at each time point was normalized to NAD(P)H intensity measurement of donor-matched quiescent fresh T cells at the 0.5-hour time point. n = 296-618 cells/condition/timepoint across 3 donors. ANOVA with three factors: donor (donor 1, 2, and 3), time (0.5-4.5 hours), and activation status (quiescent, activated). Tukey post hoc test was used to determine statistical significance for multiple comparisons between quiescent and activated groups at each time point. **(C)** Glass’s Δs to quantify effect size of activation on normalized NAD(P)H intensity of fresh and frozen T cells over time, with respect to corresponding quiescent group at each time point. **(D-G)** UMAP of 11 OMI parameters (NAD(P)H and FAD τ_m_, τ_1_, τ_2_, α_1_, α_2_, and cell size) of **(D-E)** fresh and **(F-G)** frozen T cells from 3 donors based on activation status. N =3451-4595 cells. **(H-I)** Receiver operating characteristic (ROC) curves and areas under the curve (AUCs) of Random Forest (RF) and Logistic Regression (LR) algorithms to classify **(H)** fresh and **(I)** frozen T cells by activation status (activated versus quiescent) based on OMI parameters. Data were randomly split into 70% for training (*n* = 2416-3216 cells) and 30% for testing (*n* = 1035-1379 cells). **(J)** TNF-α secreted by donor-matched fresh and frozen activated T cells after 4.5 hours of stimulation, (*n* = 3 technical replicates). Bars are mean ± SD. ** p < 0.01, *** p < 0.001, **** p < 0.0001

**Supplementary Figure 3.**
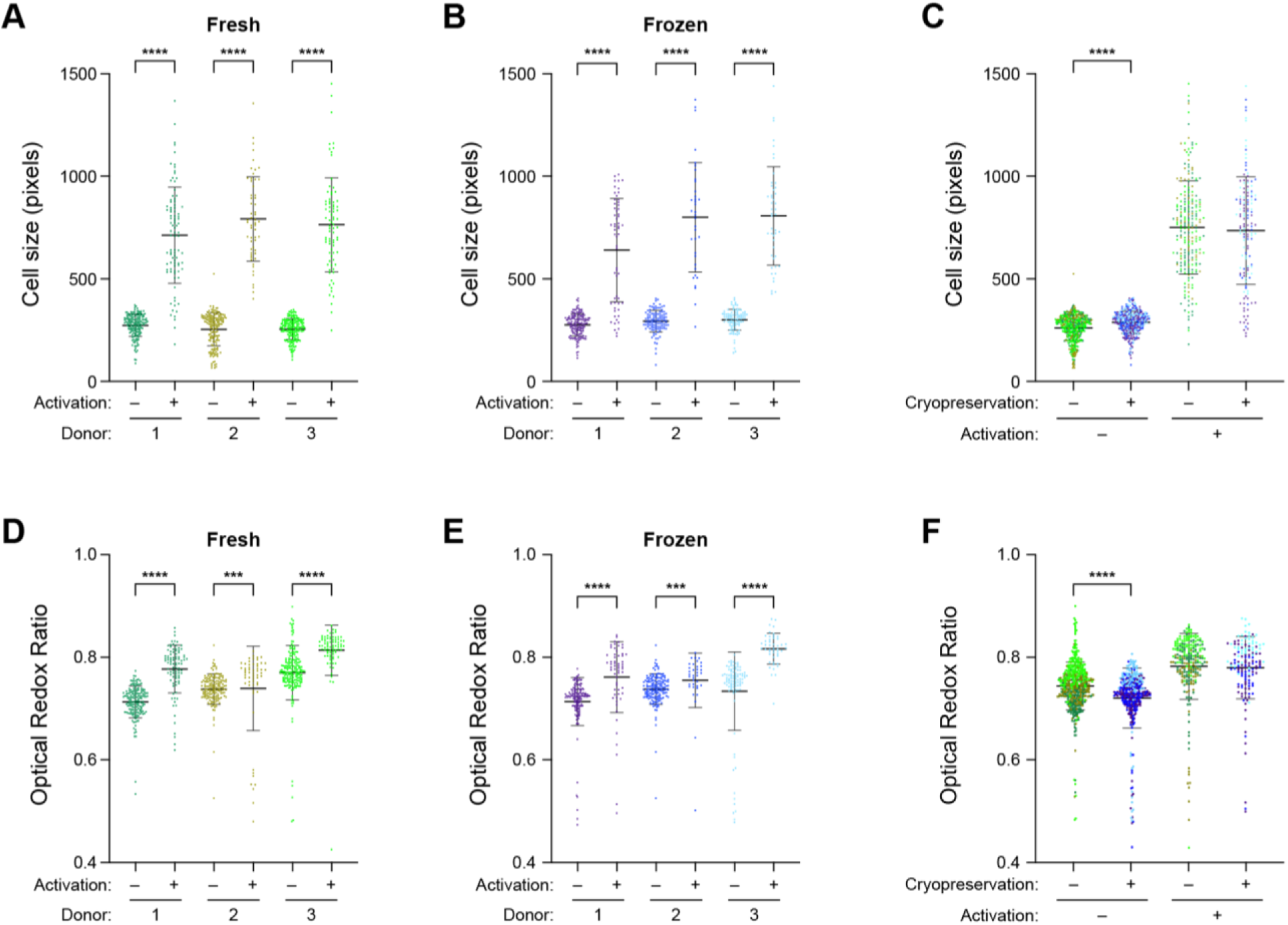
Activation response of fresh and quiescent T cells at 48 hours. **(A-B)** Quantification and **(C)** comparison of cell size of **(A)** fresh and **(B)** frozen quiescent and activated T cells at the 48-hour time point from 3 donors. **(D-E)** Quantification and **(F)** comparison of redox ratio of **(D)** fresh and **(E)** frozen quiescent and activated T cells at the 48-hour time point. For **(A-F)** n = 36-195 cells/condition/donor. Two-sided non-parametric Kruskal-Wallis test with Dunn’s post hoc tests for multiple comparisons between **(A, B, D and E)** quiescent versus activated groups for each donor and **(C, F)** fresh versus frozen for all 3 donors. *** p < 0.001, **** p < 0.0001

**Supplementary Figure 4.**
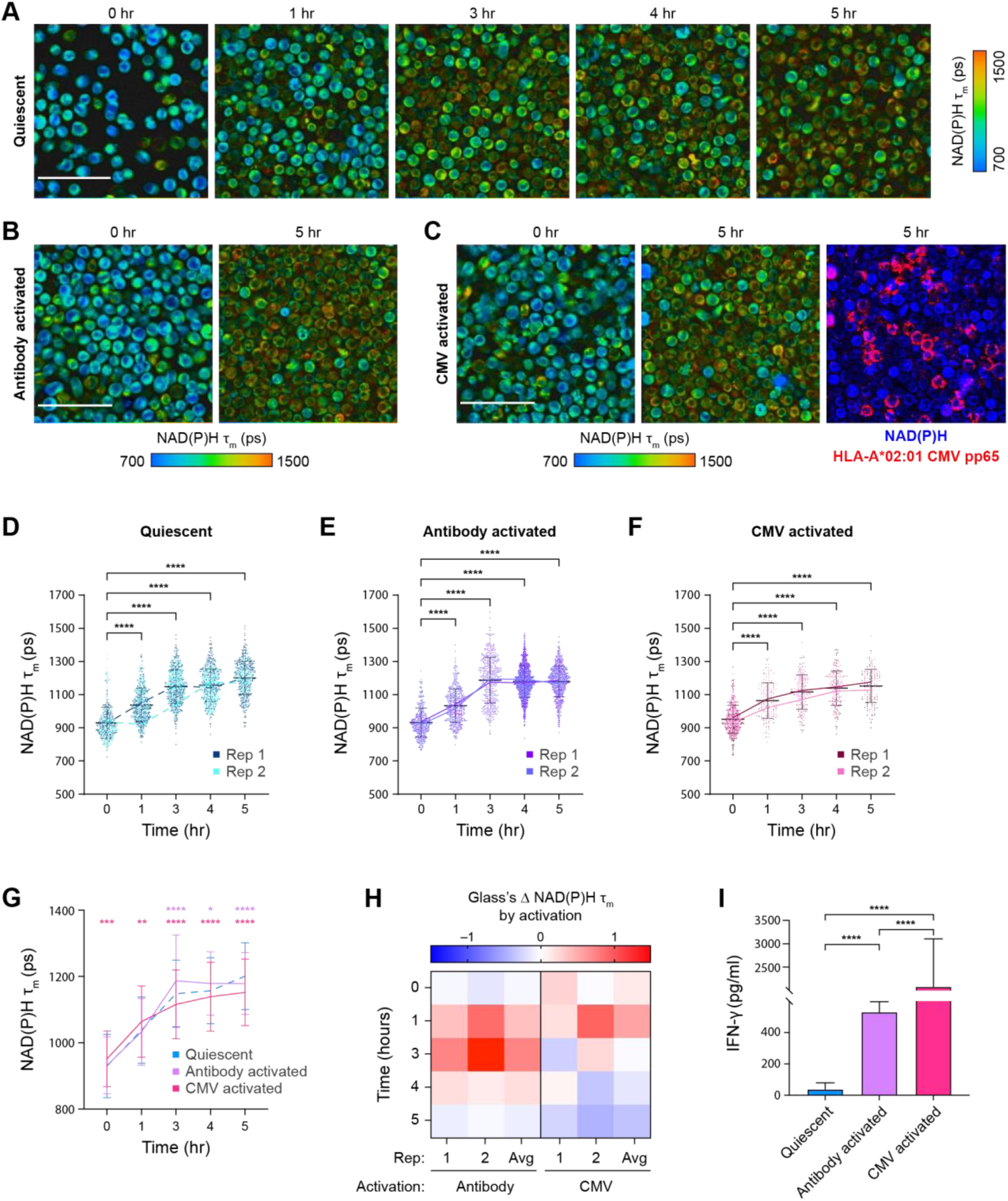
Antigen-specific activation response in frozen T cells. CMV-specific T cells were activated with two different activating stimuli: αCD2/αCD3/αCD28 antibody (antibody activation) or HLA-A*02:01 CMV pp65 peptide (CMV-peptide activation) upon thawing. **(A-C)** Representative NAD(P)H τ_m_ images of CMV-specific T cells in **(A)** quiescent, **(B)** antibody activated, or **(C)** CMV-peptide groups. APC-conjugated HLA-A*02:01 CMV pp65 peptide showed the presence of HLA-A*02:01 restricted CMV specific T cells (red). 0-hour images were collected immediately post-thaw and before activation. **(D-F)** Quantification of NAD(P)H τ_m_ of **(D)** quiescent, **(E)** antibody activated, and **(F)** CMV-peptide activated CMV-specific T cells throughout 5-hour activation time course post-thaw. Only cells stained positive for APC-conjugated HLA-A*02:01 CMV pp65 peptide (red cells in panel **(C)**) were included in the analysis for the CMV-peptide activated group. n = 84-1172 cells/condition/time point across two biologically independent CMV-specific T cell batches (replicates) from one donor. Non-parametric Kruskal-Wallis test with Dunn’s post hoc test for multiple comparisons against NAD(P)H τ_m_ measurements at the 0-hour time point. **(G)** Comparison of NAD(P)H τ_m_ between antibody activated and CMV-peptide activated T cells versus quiescent cells. Lines represent averages from two replicates of CMV-specific T cells from one donor. n = 84-1172 cells/condition/time point across 2 replicates. Two-way ANOVA with two factors: time (0, 1, 3, 4, 5 hours) and activation status (quiescent, antibody activated, and CMV-peptide activated). Dunnett’s post hoc test for multiple comparisons of activation status (antibody activated versus quiescent, and CMV-peptide activated versus quiescent) at each time point. **(H)** Glass’s Δ calculation for effect size of CD3-mediated (left) and antigen-specific (right) activation on NAD(P)H τ_m_ of cryopreserved CMV-specific T cells upon thawing. **(I)** Cytokine production by 200,000 CMV-specific T cells within 5 hours post-thaw with no stimulation (blue) or activated with αCD2/αCD3/αCD28 (purple) or HLA-matched CMV peptide (pink). n = 6 samples/condition across two batches of CMV-specific T cells. Kruskal Wallis test with Dunn’s post hoc test for multiple comparisons. Scale bar is 50μm. Bars are mean ± standard deviation. * p < 0.05, ** p < 0.01, *** p < 0.001, **** p < 0.0001.

**Supplementary Table 1.** Subject level demographic data and clinical outcomes^35^. DLBCL = diffuse large B-cell Lymphoma, MCL = mantle cell lymphoma. CR = complete response. PR = partial response. PD = progressive disease. Treatments: R=Rituximab, CHOP=cyclophosphamide, adriamycin, vincristine, prednisone, GDP=gemcitabine, dexamethasone, cisplatin, EPOCH=etoposide, prednisone, vincristine, cyclophosphamide, Adriamycin, ICE=ifosfamide, carboplatin, etoposide, BEAM=carmustine, etoposide, cytarabine, melphalan, DHAP=dexamethasone, cytarabine, cisplatin; BR=bendamustine-rituximab; MTX=methotrexate, Gem/Ox=gemcitabine, oxaliplatin.

| Patient ID | Diagnosis | Prior Lines of Treatment | Day 28 Response |
| --- | --- | --- | --- |
| 1 | DLBCL<br>MYC+<br>t(8;14) | R-CHOP, Radiation, R-GDP,<br>Rlenalidomide<br>Cell Therapy: axicabtagene ciloleucel 160<br>days prior to LV20.19 CAR | CR |
| 2 | DLBCL<br>MYC+<br>t(8;14) | R-EPOCH, R-GDP, Rlenalidomide | CR |
| 3 | DLBCL | R-CHOP, R-ICE<br>Transplant<br>BEAM auto-HCT | PR |
| 4 | MCL | R-CHOP/R-DHAP, BR,<br>acalabrutinib, R-ICE, R-EPOCH,<br>HD MTX, IT MTX, Gem/Ox<br>Transplant<br>BEAM auto-HCT | PD |

## References

1. Shah, M., Krull, A., Odonnell, L., de Lima, M. J. & Bezerra, E. Promises and challenges of a decentralized CAR T-cell manufacturing model. Front. Transplant. 2, (2023).

2. Ramakrishnan, S. et al. Should we adopt an automated de-centralized model of chimeric antigen receptor-T cells manufacturing for low-and middle-income countries? A real world perspective. Front. Oncol. 12, 1062296 (2022).

3. Whaley, D. et al. Cryopreservation: An Overview of Principles and Cell-Specific Considerations. Cell Transplant. 30, 0963689721999617 (2021).

4. de Abreu Costa, L., et al. Dimethyl Sulfoxide (DMSO) Decreases Cell Proliferation and TNF-α, IFN-γ, and IL-2 Cytokines Production in Cultures of Peripheral Blood Lymphocytes. Mol. J. Synth. Chem. Nat. Prod. Chem. 22, 1789 (2017).

5. Kondo, A. T., Kerbauy, L. N. ., Alvarez, K. C., Oliveira, D. C. ., Ribeiro, A. A. F., & Prata, K. de L. (2022). Thawing and infusion of CAR-T cell products. JOURNAL OF BONE MARROW TRANSPLANTATION AND CELLULAR THERAPY, 3(1), 165. 10.46765/2675-374X.2022v3n1p165

6. Wang, X. & Rivière, I. Clinical manufacturing of CAR T cells: foundation of a promising therapy. Mol. Ther. -Oncolytics 3, (2016).

7. Ayala Ceja, M., Khericha, M., Harris, C. M., Puig-Saus, C. & Chen, Y. Y. CAR-T cell manufacturing: Major process parameters and next-generation strategies. J. Exp. Med. 221, e20230903 (2024).

8. Pearce, E. L. Metabolism in T cell activation and differentiation. Curr. Opin. Immunol. 22, 314–320 (2010).

9. Palmer, C. S., Ostrowski, M., Balderson, B., Christian, N. & Crowe, S. M. Glucose Metabolism Regulates T Cell Activation, Differentiation, and Functions. Front. Immunol. 6, (2015).

10. Soriano-Baguet, L. & Brenner, D. Metabolism and epigenetics at the heart of T cell function. Trends Immunol. 44, 231–244 (2023).

11. Zhang, X. et al. Prolonged Cryopreservation Negatively Affects Embryo Transfer Outcomes Following the Elective Freeze-All Strategy: A Multicenter Retrospective Study. Front. Endocrinol. 12, (2021).

12. Edgar, D. H., Bourne, H., Speirs, A. L. & McBain, J. C. A quantitative analysis of the impact of cryopreservation on the implantation potential of human early cleavage stage embryos. Hum. Reprod. Oxf. Engl. 15, 175–179 (2000).

13. Gualtieri, R. et al. Mitochondrial Dysfunction and Oxidative Stress Caused by Cryopreservation in Reproductive Cells. Antioxidants 10, 337 (2021).

14. Hernández-Tapia, L. G. et al. Effects of Cryopreservation on Cell Metabolic Activity and Function of Biofabricated Structures Laden with Osteoblasts. Materials 13, 1966 (2020).

15. Ostrowska, A. et al. Investigation of Functional and Morphological Integrity of Freshly Isolated and Cryopreserved Human Hepatocytes. Cell Tissue Bank. 1, 55–68 (2000).

16. Terry, C., Dhawan, A., Mitry, R. R. & Hughes, R. D. Cryopreservation of isolated human hepatocytes for transplantation: State of the art. Cryobiology 53, 149–159 (2006).

17. Miranda, P. M. et al. Human islet mass, morphology, and survival after cryopreservation using the Edmonton protocol. Islets 5, 188–195 (2013).

18. Taylor, M. J. & Baicu, S. Review of vitreous islet cryopreservation: Some practical issues and their resolution. Organogenesis 5, 155–166 (2009).

19. Cottle, C. et al. Impact of Cryopreservation and Freeze-Thawing on Therapeutic Properties of Mesenchymal Stromal/Stem Cells and Other Common Cellular Therapeutics. Curr. Stem Cell Rep. 8, 72–92 (2022).

20. Georgakoudi, I. & Quinn, K. P. Optical imaging using endogenous contrast to assess metabolic state. Annu. Rev. Biomed. Eng. 14, 351–367 (2012).

21. Skala, M. C. et al. In vivo multiphoton microscopy of NADH and FAD redox states, fluorescence lifetimes, and cellular morphology in precancerous epithelia. Proc. Natl. Acad. Sci. 104, 19494–19499 (2007).

22. Chance, B., Legallais, V. & Schoener, B. Metabolically Linked Changes in Fluorescence Emission Spectra of Cortex of Rat Brain, Kidney and Adrenal Gland. Nature 195, 1073–1075 (1962).

23. Iyanagi, T. Molecular mechanism of metabolic NAD(P)H-dependent electron-transfer systems: The role of redox cofactors. Biochim. Biophys. Acta BBA -Bioenerg. 1860, 233–258 (2019).

24. Hu, L., Wang, N., Cardona, E. & Walsh, A. J. Fluorescence intensity and lifetime redox ratios detect metabolic perturbations in T cells. Biomed. Opt. Express 11, 5674–5688 (2020).

25. Georgakoudi, I. et al. Consensus guidelines for cellular label-free optical metabolic imaging: ensuring accuracy and reproducibility in metabolic profiling. J. Biomed. Opt. 30, S23901 (2025).

26. Kolenc, O. I. & Quinn, K. P. Evaluating Cell Metabolism Through Autofluorescence Imaging of NAD(P)H and FAD. Antioxid. Redox Signal. 30, 875–889 (2019).

27. Pham, D. L. et al. Development and characterization of phasor-based analysis for FLIM to evaluate the metabolic and epigenetic impact of HER2 inhibition on squamous cell carcinoma cultures. J. Biomed. Opt. 26, 106501 (2021).

28. Sharick, J. T. et al. Protein-bound NAD(P)H Lifetime is Sensitive to Multiple Fates of Glucose Carbon. Sci. Rep. 8, 5456 (2018).

29. Drozdowicz-Tomsia, K. et al. Multiphoton fluorescence lifetime imaging microscopy reveals free-to-bound NADH ratio changes associated with metabolic inhibition. J. Biomed. Opt. 19, 086016 (2014).

30. Kolenc, O. I. & Quinn, K. P. Evaluating Cell Metabolism Through Autofluorescence Imaging of NAD(P)H and FAD. Antioxid. Redox Signal. 30, 875–889 (2019).

31. Walsh, A. J. et al. Classification of T-cell activation via autofluorescence lifetime imaging. *Nat*. Biomed. Eng. 5, 77–88 (2021).

32. Samimi, K. et al. Time-domain single photon-excited autofluorescence lifetime for label-free detection of T cell activation. Opt. Lett. 46, 2168–2171 (2021).

33. Jones, R. G. et al. Bax and Bak regulate T cell proliferation through control of ER Ca2+ homeostasis. Immunity 27, 268–280 (2007).

34. Lemire, S. et al. Natural NADH and FAD Autofluorescence as Label-Free Biomarkers for Discriminating Subtypes and Functional States of Immune Cells. Int. J. Mol. Sci. 23, 2338 (2022).

35. Shah, N. N. et al. Bispecific anti-CD20, anti-CD19 CAR T cells for relapsed B cell malignancies: a phase 1 dose escalation and expansion trial. Nat. Med. 26, 1569–1575 (2020).

36. Dreyzin, A. et al. Immunophenotype of CAR T cells and apheresis products predicts response in CD22 CAR T cell trial for B cell acute lymphoblastic leukemia. Mol. Ther. J. Am. Soc. Gene Ther. 33, 3360–3374 (2025).

37. Pham, D. L. et al. Label-free metabolic imaging monitors the fitness of chimeric antigen receptor T cells. *Nat*. Biomed. Eng. 10.1038/s41551-025-01504–7 (2025) doi:10.1038/s41551-025-01504-7.

38. Galipeau, J. Concerns arising from MSC retrieval from cryostorage and effect on immune suppressive function and pharmaceutical usage in clinical trials. ISBT Sci. Ser. 8, 100–101 (2013).

39. Mata, M. M., Mahmood, F., Sowell, R. T. & Baum., L. L. Effects of cryopreservation on effector cells for antibody dependent cell-mediated cytotoxicity (ADCC) and natural killer cell (NK) activity in 51Cr-release and CD107a assays. J. Immunol. Methods 406, 1–9 (2014).

40. Palmisano, S. et al. Hepatocellular Metabolic Profile: Understanding Post-Thawing Metabolic Shift in Primary Hepatocytes In Vitro. Cells 14, 803 (2025).

41. Martikainen, M.-V. & Roponen, M. Cryopreservation affected the levels of immune responses of PBMCs and antigen-presenting cells. Toxicol. In Vitro 67, 104918 (2020).

42. Dreyzin, A. et al. Cryopreserved anti-CD22 and bispecific anti-CD19/22 CAR T cells are as effective as freshly infused cells. Mol. Ther. Methods Clin. Dev. 28, 51–61 (2023).

43. Keane, K. N., Calton, E. K., Cruzat, V. F., Soares, M. J. & Newsholme, P. The impact of cryopreservation on human peripheral blood leucocyte bioenergetics. Clin. Sci. 128, 723–733 (2015).

44. Jang, T. H. et al. Cryopreservation and its clinical applications. Integr. Med. Res. 6, 12–18 (2017).

45. Zhao, W., Samimi, K., Skala, M. C. & Datta, R. FLIM Playground: An interactive, end-to-end graphical user interface for analyzing single-cell fluorescence lifetime data. 2025.09.30.679625 Preprint at 10.1101/2025.09.30.679625 (2025).

